# Weakly Supervised Learning of Single-Cell Feature Embeddings

**DOI:** 10.1101/293431

**Authors:** Juan C. Caicedo, Claire McQuin, Allen Goodman, Shantanu Singh, Anne E. Carpenter

## Abstract

We study the problem of learning representations for single cells in microscopy images to discover biological relationships between their experimental conditions. Many new applications in drug discovery and functional genomics require capturing the morphology of individual cells as comprehensively as possible. Deep convolutional neural networks (CNNs) can learn powerful visual representations, but require ground truth for training; this is rarely available in biomedical profiling experiments. While we do not know which experimental treatments produce cells that look alike, we do know that cells exposed to the same experimental treatment should generally look similar. Thus, we explore training CNNs using a weakly supervised approach that uses this information for feature learning. In addition, the training stage is regularized to control for unwanted variations using mixup or RNNs. We conduct experiments on two different datasets; the proposed approach yields single-cell embeddings that are more accurate than the widely adopted classical features, and are competitive with previously proposed transfer learning approaches.

## 1. Introduction

Steady progress in automated microscopy makes it possible to study cell biology at a large scale. Experiments that were done using a single pipette before can now be replicated thousands of times using robots, targeting different experimental conditions to understand the effect of chemical compounds or the function of genes [22]. Key to the process is the automatic analysis of microscopy images, which quantifies the effect of treatments and reports relevant biological changes observed at the single-cell level. An effective computer vision system can have tremendous impact in the future of health care, including more efficient drug discovery cycles, and the design of personalized genetic treatments [3].

The automated analysis of microscopy images has been increasingly adopted by the pharmaceutical industry and thousands of biological laboratories around the world. However, the predominant approach is based on classical image processing to extract low-level features of cells. This presents two main challenges: 1) low-level features are susceptible to various types of noise and artifacts commonly present in biological experiments, being potentially biased by undesired effects and thus confounding the conclusions of a study. 2) It is unlikely that these features can detect all relevant biological properties captured by automated microscopes. With a large number of experimental conditions, the differences in cell morphology may become very subtle, and a potentially effective treatment for a disease may be missed if the vision system lacks sensitivity.

Learning representations of cells using deep neural networks may improve the quality and sensitivity of the signal measured in biological experiments. Learning representations is at the core of recent breakthroughs in computer vision [19, 26, 14], with two aspects contributing to these advances: 1) the availability of carefully annotated image databases [30, 20], and 2) the design of novel neural network architectures [15, 33]. In image-based profiling experiments, while we build on top of state-of-the-art architectures to design a solution, we face the challenge of having large image collections with very few or no ground truth annotations at all.

To overcome this challenge, we propose to train deep convolutional neural networks (CNNs) based on a weakly supervised approach that leverages the structure of biological experiments. Specifically, we use replicates of the same treatment condition as labels for learning a classification network, assuming that they ought to yield cells with similar features. Then, we discard the classifier and keep the learned features to investigate treatment similarities in a downstream analysis. This task is analogous to training a CNN for face recognition among a large number of people, and then using the learned representation to statistically identify family members in the population.

Given the high learning capacity of CNNs [36], weak labels may pose the risk of fitting experimental noise, which can bias or corrupt similarity measurements. To illustrate the problem, the feature vectors of two people might be more similar because they tend to appear in outdoor scenes more often, instead of scoring high similarity because of their facial traits. To prevent this, we evaluate two simple regularization techniques to try to control the variation of known sources of noise during the training stage.

We conduct extensive experiments with the proposed regularization strategies for training CNNs using weak labels, and evaluate the quality of the resulting features in downstream statistical analysis tasks. Our experiments were conducted on two different datasets comprising about half a million cells each, and imaged with different fluorescence techniques. The networks learned with the proposed strategies generate feature embeddings that better capture variations in cell state, as needed for downstream analysis of cell populations. The results show that our approach is an effective strategy for extracting more information from single cells than baseline approaches, yielding competitive results in a chemical screen benchmark, and improving performance in a more challenging genetic screen.

The contributions of this work are:

- A weakly supervised framework to train CNNs for learning representations of single cells in large scale biological experiments.
- The evaluation of two regularization strategies to control the learning capacity of CNNs in a weakly supervised setting. The choice of these regularizers is based on the need to control unwanted variations from batches and artifacts.
- An experimental evaluation on two datasets: a publicly available chemical screen to predict the mechanism of drugs; and a new collection of genetic perturbations with cancer associated mutations. We release the new data together with code to reproduce and extend our research.

### 2. Related Work

Weakly supervised learning is actively investigated in the vision community, given the large amounts of unlabeled data in the web. The approach has been evaluated for learning representations from Internet scale image collections [6, 17, 13] as well as for the analysis of biomedical images [35, 16]. These techniques usually deal with multi-tag noisy annotations, and adapt the loss function and training procedure to improve performance with more data. In our work, we also have noisy labels, but focus on regularization to control the variation of known sources of noise.

Given the structure of biomedical problems, various strategies have been suggested as potential solutions to the lack of labels, including transfer learning and data augmentation. An example of transfer learning is a method for diagnosing skin cancer, which reached expert-level classification performance using a CNN pretrained on ImageNet and finetuned to the specialized domain [8]. Another problem relying mostly on strong data augmentation is medical image segmentation, which was shown to work well with relatively few annotations [28].

Deep CNNs have been applied to various microscopy imaging problems to create classification models. Examples include protein localization in single cells [18] and hit selection from entire fields of view [9]. However, these are based on a supervised learning approach that requires labeled examples, and thus are limited to the detection of known phenotypes. By contrast, we aim to analyze populations of cells to explore the effects of unknown compounds or rare genetic mutations.

Recent research by Ando et al. [23] and Pawlowski et al. [25] achieved excellent results on our problem domain by transferring features from CNNs trained for generic object classification. Goldsborough et al. [10] used generative adversarial networks for learning features directly from images of cells, also bringing the ability to synthesize new images to explain variations in phenotypes. Our work is the first study to systematically train convolutional neural networks for single cells in a weakly supervised way.

#### 3. Morphological Profiling

The structure of an image collection in morphological profiling experiments is usually organized hierarchically, in batches, replicates and wells (Fig. 1). The compounds, also referred to as treatments or perturbations in this paper, are applied to cell populations, and microscopy imaging is used to observe morphological variations. Acquired images often display variations caused by factors other than the treatment itself, such as batch variations, position effects, and imaging artifacts. These factors bias the measurements taken by low-level features and are the main challenge for learning reliable representations.

**Figure 1:**
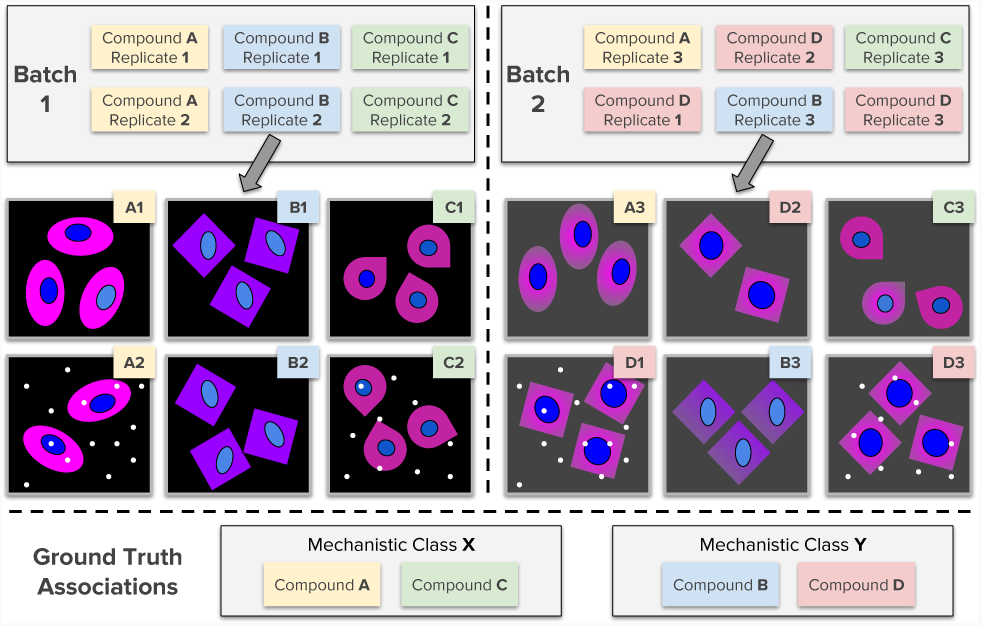
Structure of an image collection in highthroughput experiments. Compounds are applied to cells in batches and multiple replicates. Phenotypes in replicates are expected to be consistent. Note the presence of nuisance variation due to batches, microscope artifacts, or other systematic effects. The ground truth, when available, is given as known associations between compounds. The example illustrates treatments with compounds, but this could instead be genetic perturbations.

The main goal of morphological profiling is to reveal relationships among treatments. For instance, suppose a compound is being evaluated as a potential therapeutic drug and we want to understand how it affects cellular state. Depending on the morphological variations that the compound produces in cells, we may identify that it is toxic because it consistently looks similar to previously known toxic compounds. Alternatively, its morphological variations may be more correlated with the desired response for a particular disease, making it an excellent candidate for further studies. Profiling experiments may involve thousands of treatments in parallel, the majority of them with unknown response, which the experiment aims to quantify. Ground truth is usually provided as high level associations between treatments, and is very sparse (Fig. 1).

The main steps of the morphological profiling workflow include 1) image acquisition, 2) segmentation, 3) feature extraction, 4) population profiling, and 5) downstream analysis (Fig. 2). The workflow transforms images into quantitative profiles that describe aggregated statistics of cell populations. These profiles are compared using similarity measures to identify relationships. Further detail of the data analysis strategies involved in morphological profiling projects can be found in [2].

**Figure 2:**
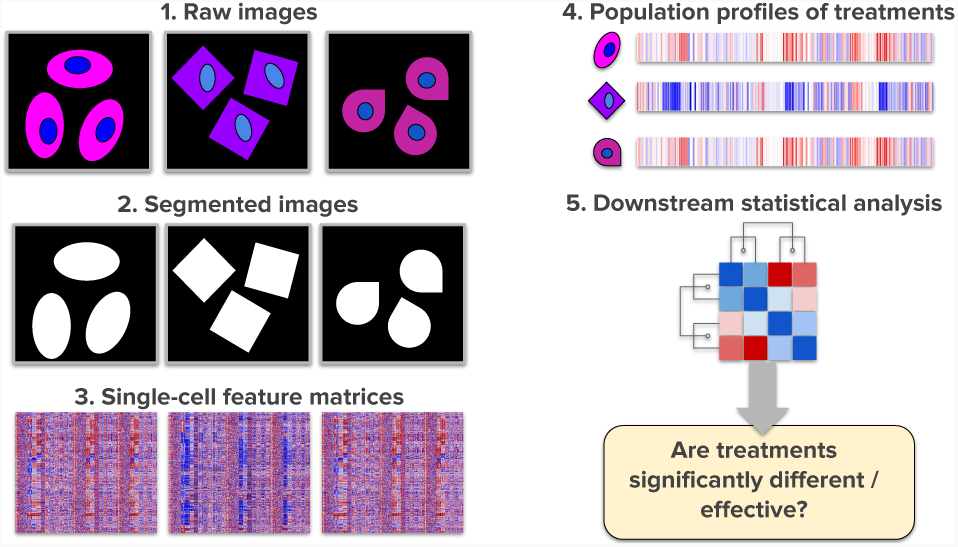
The morphological profiling workflow transforms images into quantitative measurements of cell populations. Aggregated profiles describe statistics of how one treatment altered the morphology of a population of cells. The end goal of profiling is to uncover relationships between treatments. Obtaining matrices of single-cell features in step 3 is the focus of this paper.

We focus on optimizing the features obtained in step 3 and adopt the best practices for all other steps. In particular, for image segmentation (step 2) we use the Otsu’s thresholding method in the DNA channel of the acquired image to identify single cells, which are then cropped in fixed size windows in our learning technique. We also apply the illumination correction approach proposed by Singh et al. [32] to all image channels to improve homogeneity of the field of view.

To construct the profile of one compound, we use the mean feature vector of the population of cells treated with that compound (step 4). We also apply the typical variation normalization (TVN) proposed by Ando et al. [23], using as reference distribution cells from negative control populations. The downstream statistical analysis (step 5) varies depending on the biological question. Although the discussion above has focused on treating cells with compounds (e.g. [21]), they may instead be treated with reagents that modify genes or gene products (e.g. [27]).

## 4. Learning Single-Cell Embeddings

We propose learning single-cell embeddings in a framework composed of two objectives: the *main goal* and the *auxiliary task*. The *main goal* of a morphological profiling experiment is to uncover the associations between treatments (step 5 in Fig. 2). These associations can be organized in categories or mechanistic classes as depicted in Figure 1. We define the *auxiliary task* as the process of training CNNs to recognize treatments from single cell images. The CNNs are expected to learn features that can be used in step 3 of the morphological profiling workflow.

### 4.1. Weakly Supervised Setting

Consider a collection of *n* cells *X_i_* = {*x_k_} ∀ k* ∈ {1*,…, n*}, treated with compound *Y_i_ ∈ Y*. Features can be extracted from each cell using a mapping *ϕ_θ_*(*x_k_*) ∈ ℝ^*m*^, and aggregate them using, for instance, the mean of single cell embeddings:

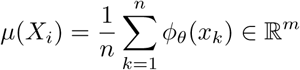

We call *µ*(*X_i_*) the mean profile of compound *Y_i_* [21]. Two compounds, *Y_i_* and *Y_j_*, have an unknown relationship *Z_i,j_* that can be approximately measured as the similarity between their mean profiles:

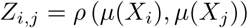

Only a few of all possible relationships *Z* between compounds (or genes) are known in biology. The problem is like identifying family members using pictures of individuals. A treatment *Y_i_* encodes the identity of an individual, and the collection of cells *X_i_* corresponds to pictures of their faces taken in different situations. Relationships such as “two individuals belonging to the same family” are represented by the unknown variables *Z_i,j_*. Our goal is to discover the true relationships *Z* using only *X* and *Y*.

We parameterize the function *ϕ_θ_*(⋅) using a convolutional neural network, whose parameters can be estimated in a data-driven manner. This CNN is trained as a multiclass classification function that takes a single cell as input and produces a categorical output. The network is trained to predict *Y* as target labels, and then the output layer is removed to compute single cell embeddings.

Notice that our *main goal* is to uncover treatment relations *Z_i,j_* instead of predicting labels *Y*, because we always know which compounds the cells were treated with. Therefore, predicting *Y* as an output is the *auxiliary task*, which we use to recovering the latent structure of morphological phenotypes in a distributed representation.

The major advantage of this approach is that it is feasible for any image-based experiment, regardless of what treatments are tested or which controls are available, and does not require knowledge of the associations between treatments. We expect the CNN to learn features that encode the high-level associations between treatments without explicitly giving it this information.

### 4.2. Noisy labels

Besides being weak labels, treatment labels are also noisy because there is no guarantee that they accurately describe the morphology of single cells. This is similar to other weakly supervised approaches that use free text attached to web images to train CNNs [6, 17].

Treatment labels are noisy for several reasons: 1) individual cells are known to not react uniformly to a treatment [12], thus generating subpopulations that may or may not be meaningful. 2) Some treatments may have no effect at all, either because they are genuinely neutral treatments or because the treatment failed for technical reasons. In either case, forcing a CNN to find differences where there are none may result in overfitting. 3) Sometimes different treatments yield the same cell phenotypes, forcing a CNN to find differences between them may, again, result in overfitting to undesired variation.

The number of replicates and single cells may be larger for some treatments than others, presenting a class imbalance during training. For this reason, we do not sample images, but instead sample class labels uniformly at random. From each sampled label we then select a random image to create training batches, following the practice in [17].

### 4.3. Baseline CNN

We first focus on training a baseline CNN with the best potential configuration to evaluate the contribution of the proposed regularization strategies. We began by evaluating several architectures including the VGG [31] and ResNet models with 18, 50 and 101 layers [15] on the auxiliary classification task. We observed similar performance in all models, and decided to choose the ResNet18 for all our experiments. This model has the least number of parameters (11.3M), helping us to limit the learning capacity as well as offering the best runtime speed.

Pixel statistics in fluorescence images are very different from those in RGB images, and we investigated several ways to normalize these values for optimal learning. The mean intensity of fluorescence images tends to be low and the distribution has a long tail, varying from channel to channel. Importantly, intensity values are believed to encode biologically relevant signal, so they are usually preserved intact or normalized to a common point of reference.

In our experiments, to our surprise, we found that preserving pixel intensities is not the best choice for CNN training. We found that rescaling pixels locally relative to the maximum intensity of each cropped cell provides the best performance. Finally, we added several data augmentation strategies, including horizontal flips, image translations, and rotations, which helped reduce the gap in performance between training and validation images.

### 4.4. RNN-based regularization

In a weakly supervised experiment, Zhuang et al. [38] noted that the association between a random image and its noisy label collected from the web is very likely to be incorrect. They suggested that looking at images in groups can reduce the probability of wrong associations, because a group of images that share the same label may have higher chance to have at least one correct example. They proposed to train CNNs that ignore irrelevant features in a group using a multiplicative gate.

We implement this concept in our work using a simple RNN architecture, specifically, using a Gated Recurrent Unit (GRU) [7] as a generic attention mechanism to track features in a group of images. The group is analyzed by the GRU as a sequence of cells in a factory line. Our recurrent architecture is many-to-one, meaning that the GRU collects information from multiple cells in a group to produce a single output for all. The GRU demands the CNN to extract relevant features from each cell for solving the classification problem (Fig. 3).

**Figure 3:**
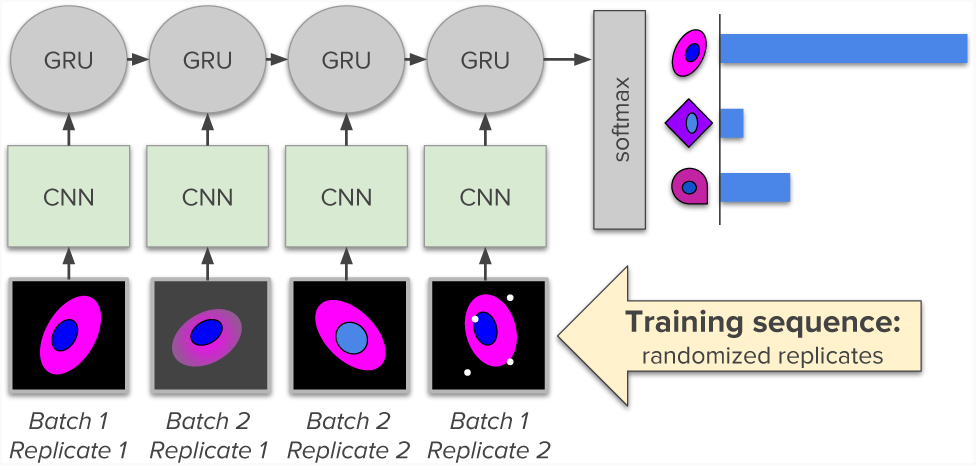
RNN-based regularization. The CNN is used as a feature extractor to inform the GRU about image contents. The GRU tracks common features in the sequence of cells to predict the correct treatment label. Cells in the sequence share the same label, have randomized order, and are sampled from different batches and replicates.

We take advantage of the sequential capabilities of a GRU in order to adapt the training process for controlling unwanted factors of variation. Our problem is not naturally sequential, allowing us to shuffle known sources of noise in a single group. First, we sample cells that share the same label in different batches and replicates. This reduces the probability of observing a group with the same technical artifact or type of noise. Second, we randomize the order of groups to prevent the GRU from learning irrelevant order-dependent patterns [34].

Importantly, the GRU trained in our models is never used for feature extraction. We leverage the ability of the recurrent network to regularize training using multiple examples at once, but after the optimization is over, we discard the RNN and only use the CNN for feature extraction. We note that similar strategies are also successfully used for training other complex systems, such as Generative Adversarial Networks (GANs) [11], in which an auxiliary discriminator network is used to train a generator.

### 4.5. Convex combinations of samples

We evaluate a regularization technique called *mixup* [37] given its ability to improve performance in classification problems that have corrupted labels, a common problem in our domain. The main idea is to artificially generate new samples by merging two data points randomly drawn from the dataset, and mixing both using a convex combination (Fig. 4). This technique has been theoretically motivated as an alternative objective function that does not minimize the empirical risk, but instead the *empirical vicinal* risk, or the distribution of inputs and outputs in the vicinity of the samples.

**Figure 4:**
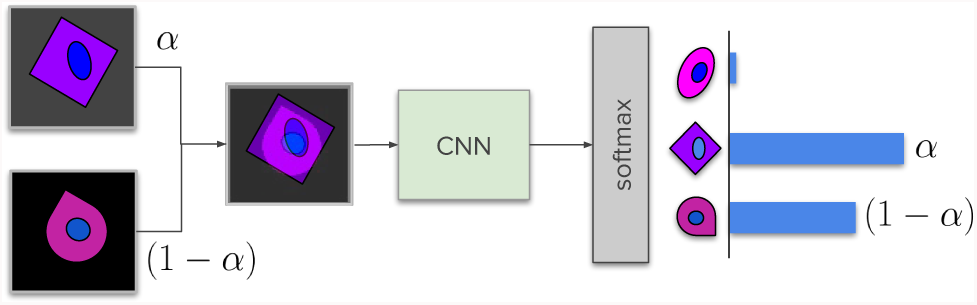
*mixup* regularization. Two cells are sampled at random from different batches and replicates, and can have different labels too. The two images are blended in a single image using a convex combination. The output is set to predict the weights of the combination.

We implement *mixup* by combining the images of two cells in a single one, weighting their pixels with a convex combination using a random parameter *α* drawn from a beta distribution. Their labels are also combined with the same weights, generating output vectors that are no longer onehot-encoded. The approach is more effective when the two sampled cells come from different replicates or batches.

*mixup* has been shown to improve performance in classification problems that have corrupted labels. In profiling datasets, both weak labels as well as input images can be corrupted. Even if the association of weak labels is correct, we face the challenge of dealing with batch effects and experimental artifacts that do not have biological meaning, but make cells look similar or different when they are not. In that sense, the images can be corrupted as well, and therefore *mixup* may help reduce the noise and highlight the right morphological features.

#### Implementation details

We create RNN models that take sequences with 3, 5 or 8 cells as input, using a GRU with 256 units. During validation, we feed sequences composed of a single cell replicated multiple times to the GRU to compare performance with the baseline. We implemented the convolutional networks used in this work in the `keras-resnet`^1^ package, and the rest of the evaluated methods are implemented in `DeepProfiler`^2^ using TensorFlow.

## 5. Experiments and Results

We present experiments on two different datasets: a study of gene mutations in lung cancer, and a publicly available benchmark of the effect of chemical compounds. Note that our experiments involve two objectives: 1) *auxiliary task*: training the CNN with weak labels using images of single cells. 2) *main goal*: analysis of population-averaged profiles to discover relationships among treatments.

### 5.1. Lung Cancer Mutations Study

Identifying the impact of mutations in cancer related genes is an ongoing research effort. In this paper, we use a collection of microscopy images that were generated to study the impact of lung cancer mutations. The imaging technique is known as *Cell Painting*, in which six fluorescent dyes are added to cells to reveal as much internal structure of the cell as possible in a single assay [1]. These are 5-channel images that can capture around 250 cells.

In this study, 65,000 images were captured, spanning about 500 gene mutations, involving more than 10 million single cells. We selected a subset of 26 variants (genes and mutations) with known impact for this work, with about

0.5 million cells. We select 10 of these variants for training CNNs, and evaluate their performance in the *auxiliary task* using a holdout set of single cells. After a network is trained, we use it to create population-averaged profiles for the *main goal* using all 26 variants (10 for training, 16 for testing).

The *main goal* of this experiment is to group variants corresponding to the same gene –regardless of mutation– together (gene accuracy), and an additional goal is to group replicates of previously unseen variants together (replicate accuracy). To this end, we use the CNN to extract features for single cells using layer conv4a (see Fig. 7) and aggregate them to the population level. Then, we analyze how often the nearest neighbors correspond to the same mutation and same gene.

**Figure 7:**
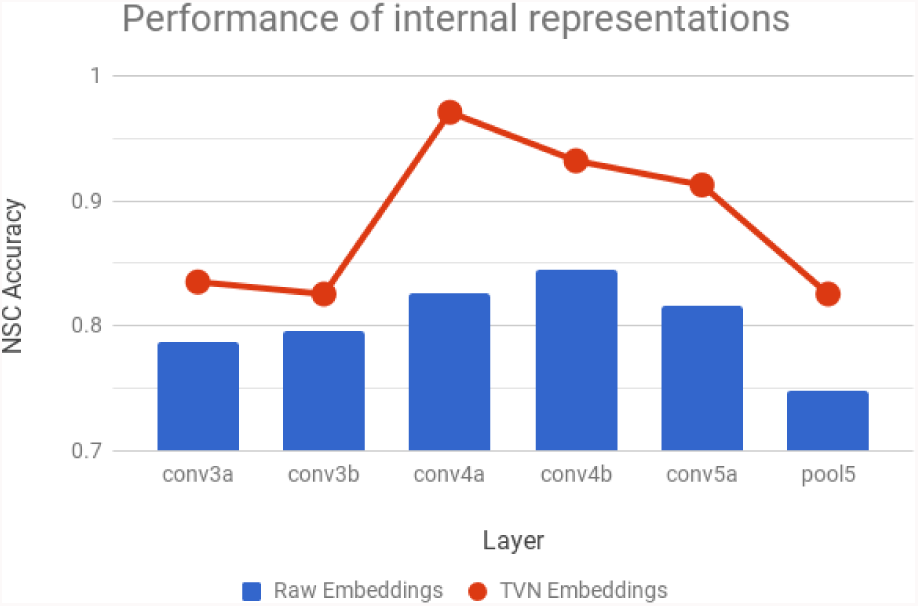
NSC classification accuracy obtained with population-averaged features extracted from different layers. This evaluation was done with a baseline CNN on the BBBC021 dataset. Intermediate layers offer better performance, while top layers specialize in solving the auxiliary task. The TVN transform [23] consistently improves performance.

Performance is improved significantly using *mixup* regularization, and only slightly using the RNN-based regularization technique (Fig 5a). The accuracy obtained with a baseline CNN in the *auxiliary task*, which is classifying single cells into 10 variants, is 70.7%. Regularization tends to decrease single-cell classification accuracy, but improves performance in the *main goal*. Gene accuracy of the 26 mutations can be improved from 69.2% to 71.6% using RNN regularization, and up to 78.3% using *mixup*.

**Figure 5:**
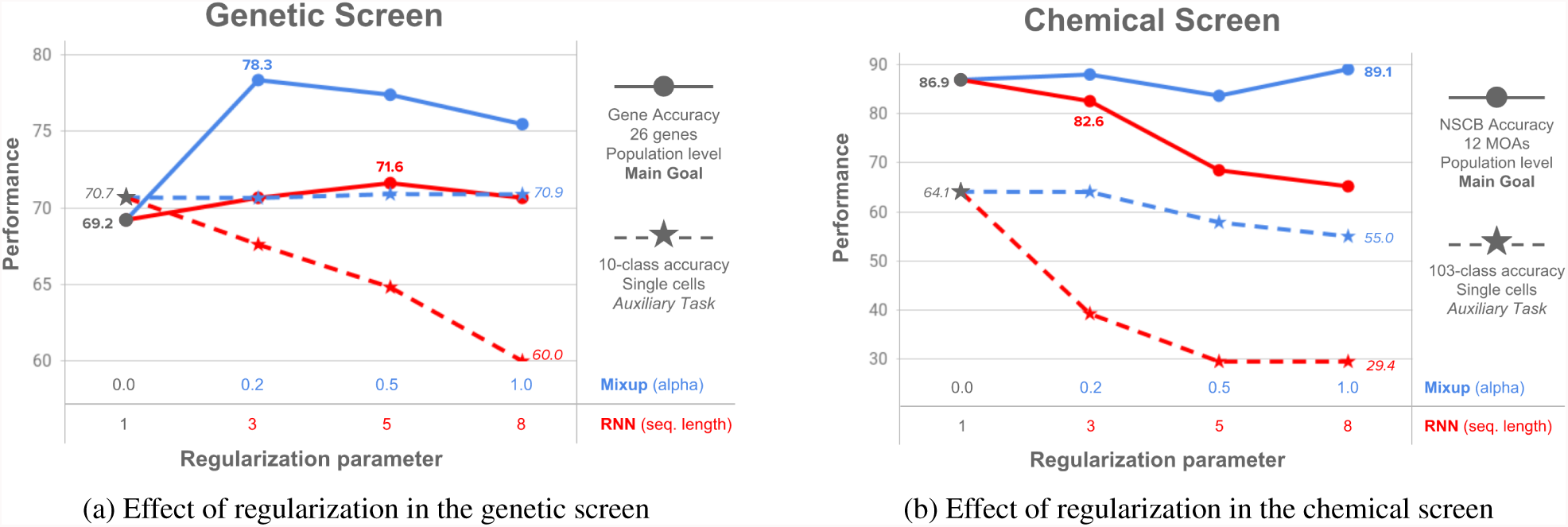
Regularization increases from left to right on the x axis, with the first parameter indicating no regularization (baseline CNN). The validation accuracy in the auxiliary task (dashed lines) tends to decrease as regularization increases. The accuracy of the main goal (solid lines) tends to improve with more regularization. a) Results in the genetic screen: both regularization techniques improve performance in the main goal, and *mixup* shows better gains. b) Results in the chemical screen: *mixup* improves slightly in the main goal, while rnn-based regularization decreases performance.

Our best CNN model yields higher replicate and gene accuracies than baseline methods, indicating that it can extract more discriminative features than previous methods (Table 1). We compare the performance to features computed by CellProfiler, our widely adopted software solution to analyze microscopy images [5], as well as features extracted by an Inception model pretrained on ImageNet as used in [25]. Figure 6 illustrates the embeddings obtained with an RNN regularized model. Notice how the classes are more clearly separated than with previous strategies, especially considering that data points are single cells.

**Table 1:** Performance of feature extraction strategies on the population-based analysis. *Replicate Accuracy* measures how often a mutation profile matches a replicate of the same mutation in the feature space. *Gene Accuracy* indicates if one mutation matches a neighbor of the same gene family.

| Model | Replicate Acc. | Gene Acc. |
| --- | --- | --- |
| ImageNet Pretrained | 42.8% | 58.2% |
| CellProfiler Features | 56.3% | 69.7% |
| Baseline CNN | 55.3% | 69.2% |
| Recurrent Model (t=5) | 58.2% | 71.6% |
| Mixup ($\alpha = 0.2$) | 59.6% | 78.4% |

**Figure 6:**
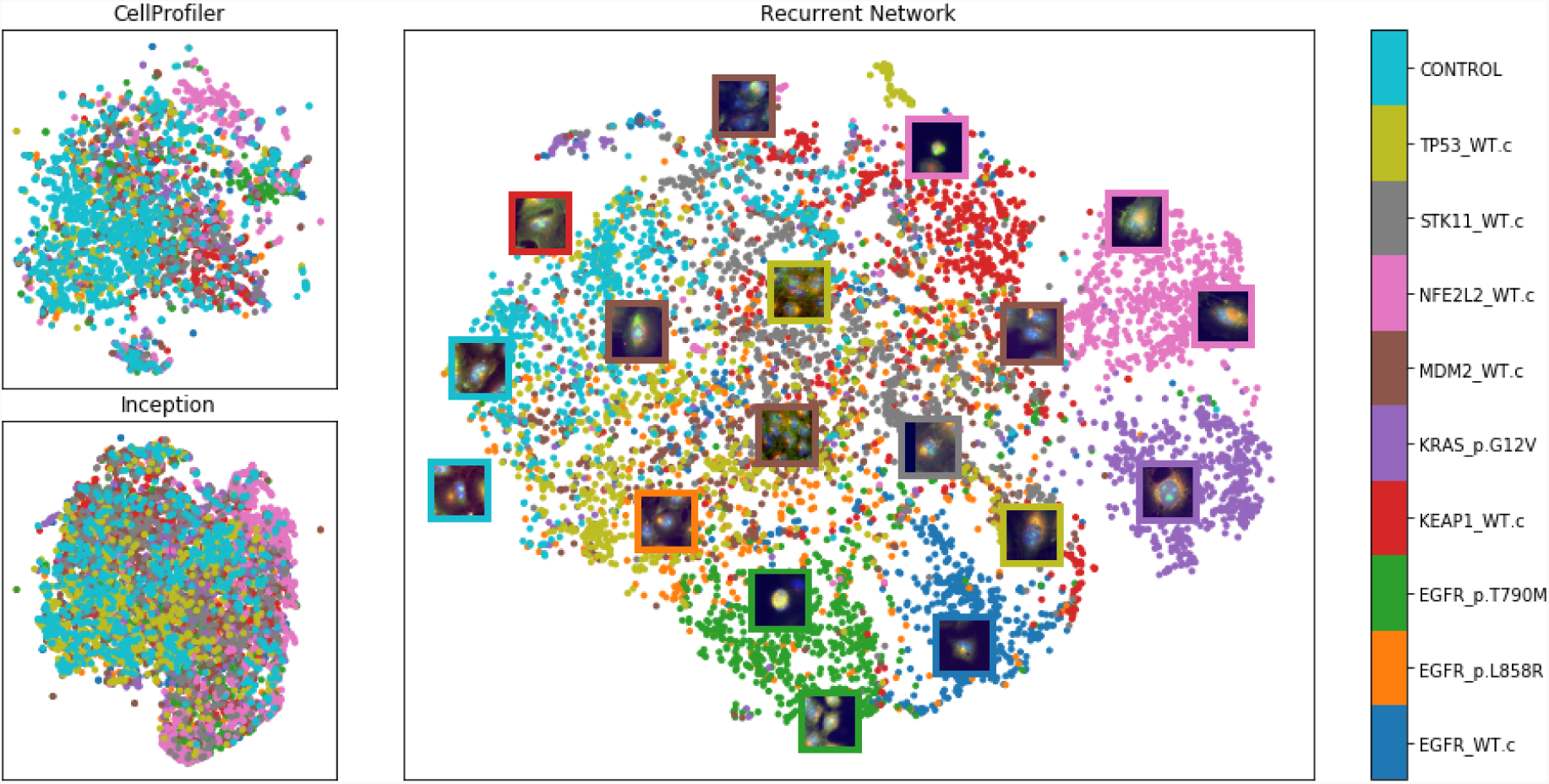
tSNE visualization of single-cell feature embeddings produced by three methods for a holdout set of the 10 training mutations. Top-left is based on classical features (with 300 dimensions). The bottom-left uses features extracted by an Inception network pretrained on ImageNet [29] (with 7,630 dimensions). The center plot shows the embeddings learned by our ResNet18 model supervised with the proposed recurrent network (256 dimensions). The colors in the plot correspond to the names of the genes used to trained the network in the auxiliary task.

### 5.2. Mechanism of Action of Compounds

The second application that we evaluate is the mechanism-of-action classification of drugs using a publicly available benchmark known as BBBC021v1 [4]. The cells in this image collection have been treated with chemical compounds of known (and strong) activity. For this reason, the relationships between compounds can be verified because drugs that are in the same mechanistic class are likely to induce similar (and detectable) responses in cells.

Images in this experiment have three channels (DNA, Actin and Tubulin), and there are a total of 35 compounds applied to cells at different concentrations. In total, 103 treatments are evaluated in this experiment, which is a subset of BBBC021v1 introduced in [21]. We choose to optimize the network to recognize these 103 treatments as the auxiliary task. In this set of experiments, we also use a ResNet18 architecture, and we assume that we have access to the full collection of batches and compounds for learning the representations of cells. This is often a valid assumption, considering that current practices optimize and normalize classical features using the samples of the same screen. We split the data in training and validation replicates to monitor performance in the auxiliary task.

After training the network in the auxiliary task, we create embeddings for cells in all treatments, and aggregate populations at the treatment level to evaluate accuracy in the main task. For this evaluation, we follow the same protocol reported in [21], which leaves one compound out to infer the class given all the others. In this way, the experiment simulates how often a new compound could be connected with the right group when its class is unknown. This evaluation is called not-same-compound (NSC) matching. We also adopt the evaluation metric proposed by Ando et al. [23] to evaluate the robustness of algorithms to batch effects and arti-facts. The simulation in this case leaves one full compound out of the analysis as well as a full experimental batch. This evaluation is called not-same-compound-and-batch (NSCB) matching, and has been shown to be more reliable.

We first note that the layer selected for creating cell embeddings is an important choice to make. Interestingly, the network disentangles the morphological structure of MOAs in an intermediate layer, before recombining these features to improve single-cell classification of treatments. Thus, layers closer to the top classification output generate features that seem to be less useful for population based analysis (Fig. 7). This evaluation was done using a baseline network without special regularization.

Applying *mixup* regularization improves performance in the *main goal*, while the RNN-based procedure tends to decrease performance (Fig. 5b). A baseline CNN can correctly classify single cells into one of the 103 treatments with 64.1% accuracy (*auxiliary task*), and obtains 86.9% accuracy in the *main goal* as measured using the NSCB procedure. We report the effect of regularization using NSCB because we expect these techniques to help reduce the impact of batch effects. From 86.9% *mixup* improves performance to 89.1% while RNNs decreases to 82.6%.

When compared to previous reports, we obtain the best NSC result using a baseline CNN, and the second best NSCB using a CNN regularized with *mixup* (Table 2). Previous strategies include classic features, transfer learning, unsupervised, and fully supervised learning. Our work is the first evaluating a weakly supervised algorithm. The transfer learning strategy using TVN proposed by Ando et al. [23] still holds the best combined result of NSC and NSCB. The difference between NSC and NSCB observed in our weakly supervised approach indicates that it is still affected by experimental artifacts and nuisance variations to some degree, despite regularization.

**Table 2:**
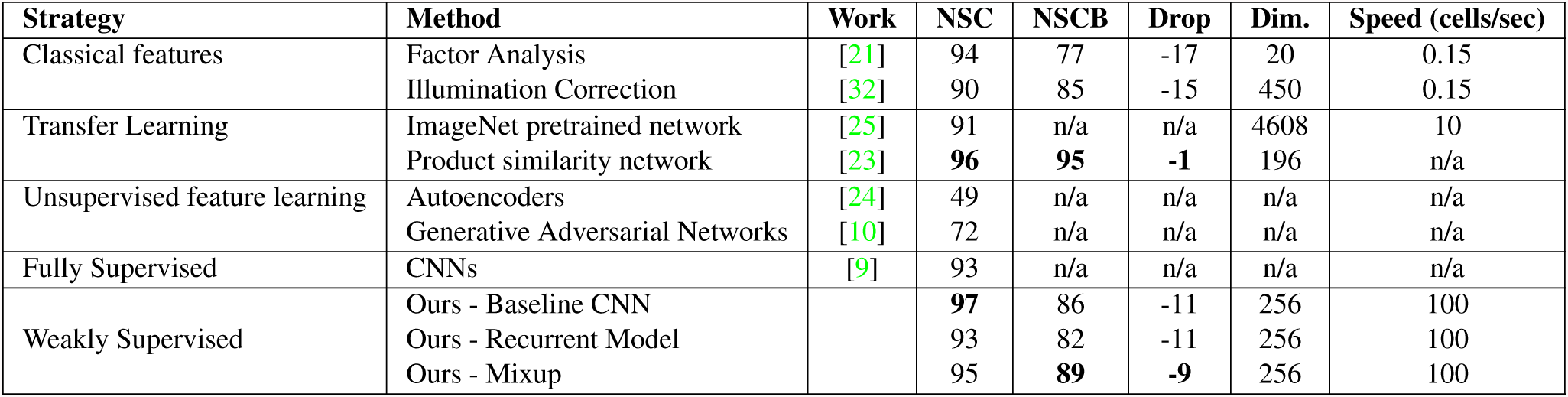
Classification accuracy on the BBBC021 chemical benchmark data set, presented alongside results of prior work. Not-Same-Compound (NSC) does not allow a match to the same compound. Not-Same-Compound-or-Batch (NSCB) does not allow a match to the same compound or any compound in the same batch. Drop is the difference between NSCB and NSC, ideally no drop in performance should be observed. Dim. refers to the dimensionality of embeddings in each layer, and Speed indicates the speed of calculating the features including Input/Output time.

Our embeddings are compact and very fast to compute with respect to transfer learning and classical features. This is in part due to GPU acceleration, and also because the only preprocessing step required to compute embeddings is cropping cells into a fixed-size window. These speeds can positively impact the cost of morphological profiling projects, requiring less computation to achieve better accuracy.

## 6. Conclusions

We described a weakly supervised learning approach for learning representations of single cells in morphological profiling experiments. A salient attribute of large scale microscopy experiments in biology is that while there is no scalable approach to annotating individual cells, there is well-structured information that can be used to effectively train modern CNN architectures using an *auxiliary task*. Our results indicate that learning representations directly from cells improves performance in the *main goal*.

The proposed methods are especially useful for studies with thousands of treatments and millions of images, in which classical features may start to saturate their discrimination ability. More research is needed to train and evaluate networks at larger scales, monitoring quality metrics that reveal performance in the absence of ground truth. The design of strategies that reduce the impact of batch effects is also an interesting future research direction, specially if these are informed by the structure of biological experiments.

## Acknowledgements

Research reported in this publication was supported by the National Institutes of Health under MIRA award number R35 GM122547. This work was also supported in part by the Chan Zuckerberg Initiative DAF, an advised fund of the Silicon Valley Community Foundation.

## Footnotes

1 https://github.com/broadinstitute/keras-resnet

2 https://github.com/jccaicedo/DeepProfiler

